# Expanded implementation of Fast & Fair paid peer review reduces time to first decision without reducing review quality in a biology journal

**DOI:** 10.64898/2026.06.02.729548

**Authors:** Daniel A. Gorelick, Alejandra Clark

## Abstract

Traditional peer review is often slowed by delays in identifying willing reviewers and waiting for completed review reports. In a 2024 pilot on the journal Biology Open, we showed that Fast & Fair peer review, which uses pre-contracted paid reviewers and a structured editorial timeline, could deliver rapid, high-quality peer. Here, we report the expanded implementation of Fast & Fair at Biology Open in 2025. From 1 April 2025 onward, all direct submissions to the journal were considered for Fast & Fair peer review unless appropriate pre-contracted reviewer expertise was unavailable. Reviewers were paid £220 per manuscript only if they completed the review on time, and the review met editorial quality expectations. Among peer-reviewed manuscripts submitted in 2025, Fast & Fair reduced time to first decision with reviews from a mean of 37.7 working days under conventional peer review to 5.5 working days. Reviewer commitment also improved. Fast & Fair invitations were accepted more often than conventional invitations (67% versus 23%), had lower nonresponse (13% versus 39%), and had higher completion among accepted invitations (98% versus 87%). Faster review was not associated with reduced review quality. Handling editors scored each review for usefulness in editorial decision-making. Fast & Fair produced fewer low-scoring reports than conventional peer review. Editorial behavior was also unchanged, with similar first-decision profiles and final acceptance rates (59% versus 61%). While financial sustainability remains to be tested at scale, the Fast & Fair model addresses a major bottleneck in traditional peer review by replacing ad hoc reviewer recruitment with conditional compensation, predefined quality standards and a strict editorial timeline.

## INTRODUCTION

Traditional peer review is a slow process, often taking weeks to identify reviewers and months before a decision following peer review is reached (Nguyen et al., 2015). Delays arise at multiple stages, but one of the most persistent bottlenecks is identifying qualified reviewers who are willing and able to provide timely, useful reports. Slow peer review delays communication of scientific findings, prolongs uncertainty for authors and editors, and can be especially burdensome for early-career researchers, trainees, and others whose career progression depends on timely publication.

Several strategies have been proposed to accelerate or improve the quality of peer review, including shorter deadlines, reminders, public recognition, reviewer certificates or credits, and monetary payments (Chan et al., 2025; Cheah and Piasecki, 2022; Chetty et al., 2014; Cotton et al., 2025; Moles et al., 2026; Yu et al., 2024; Zaharie and Seeber, 2018). Some non-monetary interventions can improve reviewer turnaround. In a field experiment at the Journal of Public Economics, reducing the review deadline from 45 to 28 days decreased median review time by 12 days (Chetty et al., 2014). In the same study, a social-incentive intervention, in which reviewers were told that their turnaround times would be publicly posted, produced a smaller improvement in review speed, particularly among senior reviewers (Chetty et al., 2014). Other forms of non-monetary recognition have been less consistently effective. Zaharie and Seeber tested whether offering public acknowledgement and reviewer certificates increased willingness to review and found that such recognition did not generally attract more reviewers; performance-contingent recognition could even reduce acceptance rates (Zaharie and Seeber, 2018). Similarly, a large quasi-natural experiment examining the Publons Global Peer Review Award found that receiving an accolade was followed by fewer subsequent reviews, suggesting that recognition-based incentives may not reliably sustain reviewer participation (Yu et al., 2024).

Monetary incentives can also affect reviewer behavior, but their effects appear to depend on the design of the incentive and the editorial workflow in which they are embedded. In the Journal of Public Economics experiment, offering reviewers $100 for submitting a report within four weeks reduced median review time by an additional 8 days (Chetty et al., 2014). More recently, a quasi-randomized trial at the journal Critical Care Medicine found that monetary incentives modestly improved review completion and reduced turnaround time by approximately 1 day without reducing review quality (Cotton et al., 2025). Some journals have also begun to implement conditional paid peer review as an operating model rather than as a short-term intervention. For example, the journal advances.in/psychology pays reviewers $100 per manuscript, with payment contingent on minimum quality requirements (Kunst, 2022). However, published data directly testing whether this operating model improves review speed or quality remains limited. Together, these studies suggest that incentives may improve peer review, but that incentive design and implementation are likely to be critical.

We previously developed the Fast & Fair peer review initiative at Biology Open to test whether rapid, high-quality peer review could be achieved in a biology journal. The initiative combines several operational changes: pre-contracted paid reviewers, explicit response and review deadlines, editor-assessed review quality standards, and a structured editorial workflow (Gorelick and Clark, 2025). In the initial 6-month pilot from July to December 2024, Fast & Fair was limited to manuscripts handled by two academic editors, while the remaining academic editors continued to handle manuscripts through conventional peer review. This design allowed us to test feasibility under controlled conditions, but it did not establish whether the model would remain effective across a wider range of editors and subject areas, or whether it would scale to the journal’s full submission volume.

Here, we evaluate the expanded implementation of Fast & Fair peer review at Biology Open. In 2025, the initiative was gradually expanded to include all academic editors. By April 1, 2025, all submissions to Biology Open, except those transferred with existing reviews, were processed through Fast & Fair peer review. Manuscripts were handled through conventional peer review only when suitable pre-contracted reviewers were not available.

We compared manuscripts handled through Fast & Fair with manuscripts handled through conventional peer review, focusing on four outcomes: time to first decision with reviews, editor-assessed review quality, editorial decision profiles and final acceptance rates, and reviewer invitation outcomes. We asked whether expanded implementation of Fast & Fair could substantially reduce review timelines without reducing review quality, altering editorial standards, or weakening reviewer commitment.

## RESULTS

### Fast & Fair reduces the time to first decision with reviews

We first asked whether the Fast & Fair workflow reduced the time from submission to first decision for manuscripts sent for peer review. We compared all peer-reviewed manuscripts at BiO between July 2024 and December 2025, excluding manuscripts rejected without peer review. Fast & Fair manuscripts reached a first decision with reviews substantially faster than manuscripts handled through conventional peer review (Figure 1). The mean time to first decision with reviews was 5.5 ± 2.2 (standard deviation) working days for Fast & Fair manuscripts compared with 37.7 ± 22.4 working days for conventional peer review. The median time to first decision was 5 working days for Fast & Fair manuscripts compared with 31 working days for conventional peer review. Because turn-around times for conventional peer review were right-skewed, we compared groups using a two-sided Mann-Whitney U test, which showed a significant reduction in review time under Fast & Fair (p<0.0001). In total, 87.1% of Fast & Fair manuscripts reached first decision with reviews within 7 working days, compared with 0% of conventionally reviewed manuscripts (the fastest conventional manuscript took 10 working days). Under Fast & Fair, the slowest manuscript took 15 working days, versus 116 working days under conventional peer review.

**Figure 1.**
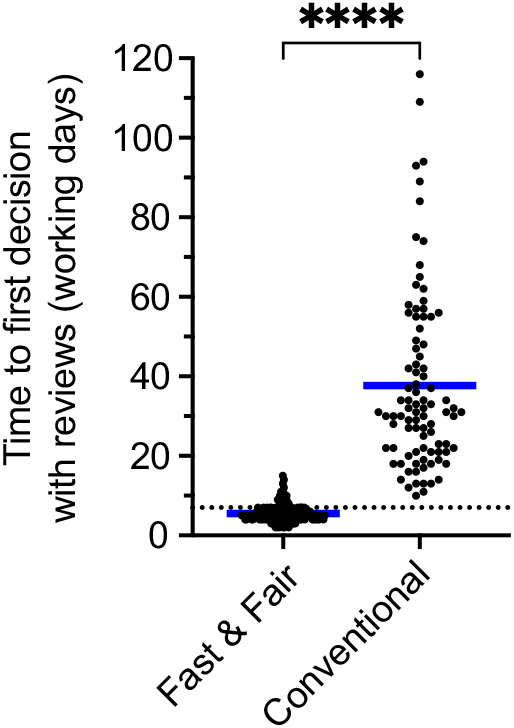
Fast & Fair peer review reduces time to first decision with reviews. Time from manuscript submission to first editorial decision after peer review for manuscripts handled through Fast & Fair or conventional peer review. Manuscripts submitted from July 2024 to December 2025. N=170 Fast & Fair manuscripts, N=91 conventional. Each data point represents one manuscript sent for peer review at Biology Open. Manuscripts rejected without peer review were excluded. Working days exclude weekends and holidays. The blue line indicates the mean for each group. The dashed line indicates the Fast & Fair target of 7 working days from submission to first decision with reviews. Groups were compared using a two-sided Mann-Whitney U test. ^****^ p<0.0001.

We also note that under Fast & Fair, the time from submission to acceptance was, on average, 64 ± 49 days (n=79 accepted manuscripts), with a median of 48 days and range of 7 to 253 days, although this largely depends on how long the authors took to address the reviewer comments and submit a revised manuscript.

### Fast & Fair shows no reduction in review quality

We next asked whether faster review was associated with reduced review quality. Handling academic editors scored each review report on a 1–100 scale, where 100 indicated that the review was helpful and appropriate for making an editorial decision, 50 indicated that the review was partially helpful, and 1 indicated that the review was not helpful for making an editorial decision. Note that, while we asked handling editors to rate each report as 1, 50 or 100 (i.e. a choice of three values with the overall scale being defined by constraints of our submission system), some editors instead rated the reviewer reports on a continuous scale from 1 to 100, complicating the interpretation of the results. Review quality scores were not lower under Fast & Fair (Figure 2). Instead, Fast & Fair review reports had slightly higher editor-assigned quality scores than conventional review reports. The mean review quality score was 97.2 for Fast & Fair reports (n=322) and 91.1 for conventional peer review reports (n=210). Median scores were 100 in both groups, reflecting the large proportion of reviews judged to be fully useful for editorial decision-making. However, low-scoring reviews were less frequent under Fast & Fair. A Mann-Whitney test showed that the distribution of review quality scores differed significantly between groups (p<0.0001). These data indicate that accelerated review under Fast & Fair did not reduce review quality and may have modestly improved the usefulness of review reports for editorial decision-making.

**Figure 2.**
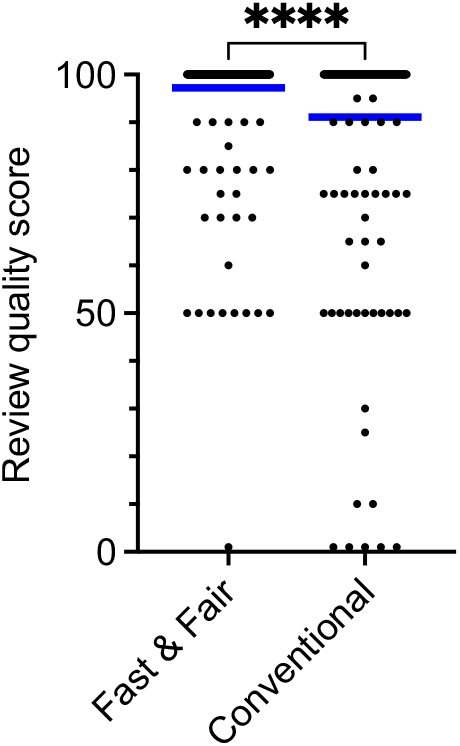
Fast & Fair peer review does not reduce review quality. Review quality scores for peer review reports submitted between July 2024 and December 2025. Each data point represents one peer review report. Handling academic editors scored each report on a 1–100 scale according to its usefulness for editorial decision-making, where 100 indicates a useful review, 50 indicates a partially useful review, and 1 indicates a review that was not useful. The blue line indicates the mean review quality score for each group. Fast & Fair: n=322 review reports, of which 294 scored 100 (91.3%). Conventional peer review: n=210 review reports, of which 167 scored 100 (79.5%). Groups were compared using a two-sided Mann-Whitney U test. ^****^ p<0.0001.

### Fast & Fair does not affect editorial decisions

We then examined whether the faster review process altered editorial decision behavior. If Fast & Fair reduced editorial rigor, this might be reflected in a different profile of decisions after peer review or in a different final acceptance rate. First, we observed that editorial rejection rates, where an editor decides to reject a manuscript without peer review, remained similar (~37% of all submitted manuscripts) in 2025 compared to previous years where most or all manuscript went through conventional peer review (Gorelick, 2026). Next, we compared first decisions following peer review and final outcomes for peer-reviewed manuscripts handled through Fast & Fair or conventional peer review between July 2024 and December 2025 (Table 1). The distribution of first decisions after peer review was similar between groups. Among Fast & Fair manuscripts, 52 of 170 were rejected (31%), 34 received minor revision decisions (20%), and 84 received major revision decisions (49%). Among conventionally reviewed manuscripts, 29 were rejected (32%), 20 received minor revision decisions (22%), and 40 received major revision decisions (44%). The distribution of first decisions did not differ between groups (χ^2^=0.49, df=2, p=0.78). Final acceptance rates were also similar: 79 of 135 Fast & Fair manuscripts with final outcomes were accepted (59%), compared with 50 of 82 conventional manuscripts (61%; Fisher’s exact test, p=0.78). Thus, the faster review timeline did not measurably alter editorial decision-making. We conclude that editorial decisions were not influenced by financial considerations.

**Table 1.** Editorial decisions and final outcomes for peer-reviewed manuscripts handled through Fast & Fair or conventional peer review. Manuscripts sent for peer review between July 2024 and December 2025 were included; manuscripts rejected without peer review were excluded. First-decision outcomes are shown as the number and percentage of manuscripts receiving a reject, minor revision, major revision or accept decision after peer review. For first decisions, n denotes the total number of manuscripts sent for peer review in each workflow. Final acceptance rates are shown separately using the subset of manuscripts for which a final editorial outcome had been reached by the time of analysis. For final outcomes, n denotes the total number of manuscripts with a final decision. Decisions following peer review (reject, minor revision, major revision) did not differ between Fast & Fair and conventional peer review (χ^2^=0.49, df=2, p=0.78), and final acceptance rates were similar between groups (59% vs 61%; Fisher’s exact test, p=0.78). Note that two manuscripts were accepted following peer review, without additional revisions, because they were submitted with reviews from another journal.

|  | Fast & Fair<br>n (%) | Conventional<br>n (%) |
| --- | --- | --- |
| Total manuscripts sent for peer review | 170 | 91 |
| Reject | 52 (31%) | 29 (32%) |
| Minor revision | 34 (20%) | 20 (22%) |
| Major revision | 84 (49%) | 40 (44%) |
| Accept | 0 (0%) | 2 (2%) |
| Total manuscripts where a final decision was made | 135 | 82 |
| Final decision = accept | 79 (59%) | 50 (61%) |

### Fast & Fair improves reviewer commitment

Finally, we asked whether Fast & Fair affected reviewer commitment. Because retainer reviewers in the 2024 pilot were contractually required to accept review invitations, this analysis was limited to calendar year 2025, when Fast & Fair used the freelance reviewer model. The unit of analysis was the review invitation, not the unique reviewer. Reviewer invitation outcomes were classified as accepted, declined, or no response. For conventional peer review, no response was defined as no reply within 10 total days. For Fast & Fair, no response was defined as no reply within 1 working day.

Fast & Fair substantially improved reviewer responsiveness and commitment (Table 2). In 2025, 307 of 457 Fast & Fair review invitations were accepted (67%), compared with 79 of 347 conventional peer review invitations (23%). Nonresponse was much lower under Fast & Fair: 59 of 457 Fast & Fair invitations received no response (13%), compared with 137 of 347 conventional review invitations (39%). Declines were also less frequent under Fast & Fair than conventional peer review (21% versus 37%). A 2×3 chi-square test comparing invitation outcomes showed a significant difference between Fast & Fair and conventional peer review (χ^2^=156.9, df=2, p<0.0001).

**Table 2.** Reviewer invitation outcomes for Fast & Fair and conventional peer review in 2025. Reviewer commitment was assessed using all reviewer invitations sent during calendar year 2025. Outcomes were classified as accepted the invitation to review, declined, or did not respond. “Completed” indicates that a reviewer who accepted an invitation submitted a review report; completion percentages are calculated using accepted invitations as the denominator. Reviewer invitation outcomes differed between Fast & Fair and conventional peer review (χ^2^ test comparing accepted, declined, and did not respond; p<0.0001). Among reviewers who accepted invitations to review, review completion was also higher under Fast & Fair than conventional peer review (Fisher’s exact test; p<0.0001).

|  | Fast & Fair<br>n (%) | Conventional<br>n (%) |
| --- | --- | --- |
| Total manuscripts | 150 | 39 |
| Total reviewers invited | 457 | 347 |
| Accepted invitation to review | 307 (67%) | 79 (23%) |
| Completed the review | 302 (98%) | 69 (87%) |
| Declined invitation to review | 94 (21%) | 127 (37%) |
| Did not respond | 59 (13%) | 137 (39%) |

Among reviewers who accepted invitations, review completion was also higher under Fast & Fair. Fast & Fair reviewers submitted reports for 302 of 307 accepted invitations (98%), compared with 69 of 79 accepted invitations under conventional peer review (87%). Fisher’s exact test showed that completion among accepted invitations was significantly higher under Fast & Fair (p<0.0001). Among reviewers who accepted invitations, 6 review reports (2%), were overdue: 3 were submitted one day late, and the other 3 were never submitted (after 1 day late, we solicited additional reviews so as not to delay the manuscript). These data indicate that Fast & Fair increased reviewer commitment at multiple stages: reviewers were more likely to accept invitations, less likely not to respond, and more likely to complete the review after accepting. These outcomes should be interpreted considering the structure of the peer review contract: Fast & Fair reviewers were not paid merely for accepting invitations, but only for submitting timely reports that met editorial quality expectations.

### The costs of Fast & Fair peer review

The 2025 expansion of Fast & Fair required direct reviewer payments and additional staff support to operate the workflow at larger scale (Table 3). Reviewer payments totaled approximately £66,000 in 2025. Additional staff costs for Fast & Fair were estimated at £30,000–£40,000 and included time from the journal managing editor, administrators, and technical support for system optimisation, monitoring the initiative, and reviewer contract management. Optimisation of the Fast & Fair contract management tool added £8,000. The total estimated cost of the 2025 Fast & Fair reviewer initiative was therefore £104,000–£114,000.

**Table 3.** Estimated costs of the Fast & Fair reviewer initiative in 2025. Reviewer payments include payments to freelance reviewers for completed reviews that met time and quality expectations. Staff costs include estimated costs for system optimisation, monitoring the initiative, and reviewer contract management. Staff = journal managing editor, administrators and technical support.

| Fast & Fair reviewer initiative 2025 | Cost (£) |
| --- | --- |
| Payments to reviewers | 66,000 |
| Staff costs | 30,000-40,000 |
| Contract management tool optimisation | 8,000 |
| Total estimated | 104,000-114,000 |

The costs of Fast & Fair peer review decreased substantially on a per-manuscript basis as the initiative expanded. During the 2024 pilot, 20 Fast & Fair manuscripts were peer reviewed at an estimated total cost of £50,635–£60,635, corresponding to £2,532–£3,032 per manuscript (Gorelick and Clark, 2025). In 2025, 150 Fast & Fair manuscripts were peer reviewed at an estimated total cost of

£104,000–£114,000 (Table 3), corresponding to £693–£760 per manuscript. Thus, we were able to expand Fast & Fair while reducing the estimated cost per manuscript by approximately 74% (based on the midpoint of each cost range). Several reasons contribute to the reduction in costs: fixed costs remained steady or decreased (eg the fixed costs of setting up the reviewer contracts); the amount of time staff devoted to running Fast & Fair is less than the time required to set up Fast & Fair; shifting from the retainer model (in 2024) to the freelance model (in 2025) of paying peer reviewers. This decrease in the cost per manuscript is notable because The Company of Biologists is a small nonprofit publisher and therefore benefits less from economies of scale compared with larger commercial publishers. Continued expansion, workflow optimization, and appropriate adjustment of article processing charges should further improve the financial sustainability of Fast & Fair peer review.

## DISCUSSION

This expanded implementation of Fast & Fair peer review substantially reduced the time that manuscripts spent under review at Biology Open. Manuscripts reviewed through Fast & Fair reached a first decision with reviews in a median of 5 working days, compared with 31 working days for manuscripts handled through conventional peer review. This reduction was achieved while maintaining review quality, preserving editorial decision profiles, and improving reviewer responsiveness. A major advantage of Fast & Fair was the reduction in reviewer-invitation burden. On average, Biology Open invited 3.05 reviewers per Fast & Fair manuscript compared with 8.90 reviewers per conventionally reviewed manuscript, meaning that conventional peer review required nearly three times as many reviewer invitations per manuscript. Together, these findings extend our initial pilot study by showing that rapid peer review can be implemented beyond a small, two-editor feasibility experiment and maintained across a broader journal workflow.

The most direct effect of Fast & Fair was the reduction in time to first decision with reviews. Conventional peer review contains several points at which manuscripts can stall: identifying appropriate reviewers, waiting for reviewers to respond, inviting additional reviewers after declines or nonresponses, waiting for reports, and then returning the manuscript to the handling editor for a decision. Fast & Fair addresses these delays by replacing an ad hoc reviewer search process with a structured workflow involving pre-contracted reviewers, explicit response expectations, explicit review deadlines, and editor oversight. The effect size was large. Fast & Fair manuscripts were reviewed in approximately one-sixth of the time required for conventional peer review. This suggests that much of the delay in conventional peer review at a sound science journal is not intrinsic to the intellectual work of evaluating a manuscript but instead arises from the organization of the review process.

The time from submission to acceptance was reduced to 48 total days (median) under Fast & Fair. While much of the time from submission to acceptance is out of our control because authors can take months to submit a revised manuscript, it is likely that the speed of the initial peer review contributed to the reduction in total time to acceptance.

A central concern with rapid peer review is that speed could come at the expense of rigor. Our data do not support that concern. Editor-assessed review quality was not lower under Fast & Fair. Instead, review quality scores were modestly higher for Fast & Fair reports than for conventional reports, with fewer low-scoring reviews. This does not mean that Fast & Fair necessarily causes reviewers to write better reviews, but it does indicate that shorter deadlines and financial compensation did not reduce the usefulness of reports for editorial decision-making. One possible explanation is that the Fast & Fair system selects for reviewers who are willing to commit to a defined task within a defined time frame. Another possibility is that the contract structure, including explicit expectations for both timeliness and quality, encourages reviewers to take the assignment seriously. In either case, the data argue against the assumption that fast review leads to superficial review.

Fast & Fair also did not measurably alter editorial standards. If faster peer review compromised editorial rigor, one might expect a lower rejection rate after peer review, a shift toward less demanding revision decisions, or a higher final acceptance rate. We did not observe these patterns. First-decision profiles after peer review were similar between Fast & Fair and conventional peer review, and final acceptance rates were similar among manuscripts with final outcomes. These findings suggest that Fast & Fair accelerated the process by which editorial decisions were reached rather than changing the decisions themselves.

The reviewer invitation data provide insight into why Fast & Fair reduced review times so substantially. Reviewers invited through Fast & Fair were more likely to accept invitations, less likely not to respond, and more likely to complete reviews after accepting. This improvement in reviewer commitment is probably central to the success of the model. In conventional peer review, reviewer nonresponse is a major source of delay because editors often wait days or weeks before inviting additional reviewers. In Fast & Fair, reviewers are asked to make a rapid decision about availability, and the system is designed around that commitment. Fast & Fair did not simply pay reviewers for agreeing to review. Reviewers were paid £220 only if they submitted the report within the required timeframe, and the report met editorial quality expectations. This conditional compensation structure likely contributed to the higher acceptance, lower nonresponse, and higher completion rates observed under Fast & Fair.

Fast & Fair may also have implications for equity in peer review, although this study did not directly test that possibility. Recent work analyzing biomedical and life science articles indexed in PubMed found that female-authored articles spent longer under review than male-authored articles, with median review times 7.4% to 14.6% longer depending on author gender, and also found longer review times for authors based in low-income countries (Alvarez-Ponce et al., 2026). Fast & Fair should not be assumed to eliminate bias. However, by standardizing and compressing the review timeline, it may reduce the procedural space in which delay-based disparities arise. The same response expectations, review deadlines, and compensation structure apply regardless of author identity, institution, geography, or career stage. Even if residual differences in review time persisted, their practical magnitude may be smaller within a workflow where most manuscripts receive decisions within days rather than weeks or months. Future studies should test this directly by examining whether Fast & Fair reduces disparities in review time by author gender, geography, institutional setting, or career stage.

The present results also raise questions about scalability. Our data support expanded implementation within a single biology journal, but they do not prove that Fast & Fair can be generalized across all journals, disciplines, or publishing models. Scalability depends on maintaining a sufficiently large and diverse pool of pre-contracted reviewers, matching reviewer expertise to manuscript content, preserving editor engagement, and ensuring financial sustainability. As Fast & Fair expands, the main bottleneck may shift from finding willing reviewers to maintaining an adequate reviewer pool across subject areas. Some manuscripts still require conventional peer review when suitable pre-contracted reviewers are unavailable. This indicates that reviewer coverage, not only reviewer speed, is a critical determinant of scalability.

Several limitations should be noted. First, this was not a randomized trial. Manuscripts were not randomly assigned to Fast & Fair or conventional peer review, and manuscripts handled conventionally may differ from Fast & Fair manuscripts in ways that influence review time or editorial outcome. Some manuscripts were handled through conventional review because the journal lacked appropriate pre-contracted reviewer expertise. Second, this study was conducted at a single journal. Biology Open has a particular editorial model, scope, staffing structure, and relationship with The Company of Biologists, and the results may not generalize to journals with different workflows, submission volumes, editorial policies, or financial constraints. Third, although the sample size is substantially larger than in the initial pilot, it remains modest relative to the scale of peer review across biomedical publishing.

Fourth, review quality was assessed by handling academic editors rather than by an independent blinded panel. This was appropriate for the operational question we asked, because the relevant issue was whether reviews were useful for editorial decision-making. However, editor-assessed quality may reflect editor expectations, manuscript complexity, or variation among editors. For accepted manuscripts, we publish the review reports, which are freely available for assessing review quality. Fifth, the long-term effects of financial incentives remain unknown. Fast & Fair could improve reviewer commitment in the short term but still be vulnerable to reviewer fatigue or motivation crowding over longer periods. Continued monitoring of reviewer retention, repeat participation, and review quality will be essential.

Finally, this report does not address the financial sustainability of the model, although this remains an active area of analysis. Fast & Fair requires payment to reviewers and administrative costs for recruiting reviewers, managing contracts, monitoring timelines, and supporting editors. A key challenge is that these costs are incurred whether or not a manuscript is accepted and generates article processing charge (APC) revenue, while editorial decisions must continue to remain independent of financial considerations. We are currently evaluating how modest APC increases could offset these costs without placing undue burden on authors, noting that fee waivers are available where needed. An important test will be whether authors and funding agencies value faster review sufficiently to support a higher APC. Ultimately, sustainability depends on submission volume, reviewer costs, acceptance rates after peer review, and the extent to which operational efficiencies can reduce expenses. These financial constraints may be especially important for nonprofit publishers and smaller journals that lack the economies of scale of large commercial publishers.

In summary, expanded implementation of Fast & Fair peer review markedly reduced the time to first decision with reviews, without reducing editor-assessed review quality or altering editorial decision outcomes. Fast & Fair also improved reviewer responsiveness and completion, suggesting that explicit expectations paired with compensation can strengthen reviewer commitment. These findings support Fast & Fair as a viable alternative to conventional peer review in a biology journal and justify further testing across additional journals, disciplines, and reviewer-recruitment models.

## MATERIALS AND METHODS

### Study design and implementation

This study evaluated the expanded implementation of Fast & Fair peer review at Biology Open. During the initial pilot from July to December 2024, Fast & Fair was limited to manuscripts handled by two academic editors, while manuscripts handled by the remaining academic editors continued through conventional peer review. During 2025, Fast & Fair was progressively expanded to all academic editors. By April 1, 2025, all manuscripts submitted to Biology Open were considered for Fast & Fair peer review unless the journal lacked pre-contracted reviewers with the appropriate subject-matter expertise, in which case manuscripts were handled through conventional peer review.

Fast & Fair manuscripts followed a structured editorial workflow. After submission, editorial staff assessed manuscripts for compliance with Biology Open’s publication criteria, including scientific soundness and potential indicators of paper mill activity. Manuscripts that passed this bar were sent to a handling academic editor, who decided whether to reject the paper (based on obvious problems according to our <u>peer review rubric</u>) or to send for peer review. The handling academic editor invited two peer reviewers. Fast & Fair reviewers were pre-contracted and were expected to respond to invitations within one working day and, if they accepted, submit their review reports within four working days. After both reviews were received, the handling editor assessed the reports and issued a first decision.

Conventional peer review followed Biology Open’s standard editorial workflow. Under conventional review, academic editors identified and invited reviewers through the usual process, and reviewers were requested to provide a review in 10 days, though this is not enforced. Editors handling conventional manuscripts were instructed not to use the Fast & Fair pool of pre-contracted reviewers.

### Reviewer contracts and compensation

Fast & Fair reviewers were recruited and contracted before manuscript assignment. From July through December, 2024, we signed reviewers to either freelancer contracts (payment of 220 GBP per manuscript) or retainer contracts (fixed payment of 600 GBP per quarter) (Gorelick and Clark, 2025). From 1 January 2025, all reviewers under the retainer contract were transferred to freelancer contracts. On 25 March 2025, an open call was launched inviting eligible reviewers to apply for freelancer contracts: https://www.biologists.com/bio-fast-and-fair-peer-review/. Reviewers were paid £220 per manuscript. Payment was conditional on meeting both timeliness and quality expectations. Reviewers were required to accept or decline an invitation within one working day and, if they accepted, to submit the review within four working days. Reviewers were paid only if the report was submitted on time and judged by handling editors to meet the journal’s quality expectations. Review quality was assessed based on whether the report was useful for editorial decision-making, as described below (Review Quality). Reports that were late or not helpful for editorial decision-making (score 1) were not eligible for payment. By 1 May 2026, the Fast & Fair reviewer pool included approximately 420 pre-contracted freelance reviewers across the subject areas covered by Biology Open.

This contract structure was designed to link compensation to performance rather than to review invitation. Thus, Fast & Fair differed from conventional peer review not only by paying reviewers, but by combining payment with pre-commitment, explicit deadlines, quality standards, and editorial oversight.

### Time to first decision with reviews

To determine whether Fast & Fair reduced review timelines, we compared the time from manuscript submission to first decision with reviews for manuscripts handled through Fast & Fair or conventional peer review. The analysis included manuscripts submitted to Biology Open that were sent for peer review. Manuscripts rejected without peer review were excluded. Transferred manuscripts that arrived with existing peer review reports from another journal were excluded from Fast & Fair, because these manuscripts did not undergo a full Biology Open peer review process. In the cases where an editor determined that a manuscript transferred from another journal with reviews required additional peer review reports, these were handled via the conventional peer review process.

Time to first decision with reviews was measured in working days from submission to the date of the first editorial decision after peer review. Working days excluded weekends and public UK holidays when the editorial office is closed. Manuscripts were included regardless of subject area. Each data point represents a single peer-reviewed manuscript. The blue line in the figure indicates the mean for each group.

Because conventional peer review turnaround times were right-skewed and not normally distributed, Fast & Fair and conventional peer review were compared using a two-sided Mann-Whitney U test. Mean, median, interquartile range, range, and the proportion of manuscripts reaching first decision within seven working days were calculated for each group.

### Review quality

To assess whether faster peer review affected review quality, handling academic editors scored each peer review report for its usefulness in editorial decision-making. Scores were assigned on a continuous 1–100 scale. A score of 100 indicated that the review was helpful for making an editorial decision, a score of 50 indicated that the review was partially helpful, and a score of 1 indicated that the review was not helpful. In addition, to ensure that the fast turnaround times of the Fast & Fair editorial workflow did not compromise the quality of editorial assessment, all manuscripts received internal oversight from editorial staff.

The analysis included review reports submitted between July 2024 and December 2025 for manuscripts handled through Fast & Fair or conventional peer review. The unit of analysis was the individual peer review report, not the manuscript. Because the primary goal was to compare review reports as experienced by editors, review quality was analyzed at the report level. Each data point represents a single review report.

Because review quality scores were bounded from 1 to 100 and clustered near the upper end of the scale, scores were compared between Fast & Fair and conventional peer review using a two-sided Mann-Whitney U test. Mean, median, interquartile range, and range were calculated for each group.

### Editorial decisions and final outcomes

To determine whether Fast & Fair affected editorial decision-making, we compared first decisions after peer review and final outcomes between manuscripts handled through Fast & Fair or conventional peer review. This analysis included manuscripts sent for peer review between July 2024 and December 2025. Manuscripts rejected without peer review were excluded.

First decisions after peer review were categorized as reject, minor revision, or major revision. The unit of analysis was the manuscript. The distribution of first-decision categories was compared between Fast & Fair and conventional peer review using a 2×3 chi-square test.

Final outcomes were analyzed separately because not all manuscripts had reached a final decision by the time of analysis. For the final-outcome analysis, the denominator was the number of peer-reviewed manuscripts for which a final decision had been made. Final acceptance rate was calculated as the number of accepted manuscripts divided by the number of manuscripts with a final outcome in each group. Final acceptance rates were compared between Fast & Fair and conventional peer review using Fisher’s exact test.

### Reviewer commitment

To assess reviewer commitment, we analyzed reviewer invitation outcomes during calendar year 2025. The 2024 pilot period was excluded from this analysis because retainer reviewers used during the pilot were contractually required to accept review invitations and therefore were not comparable to conventional reviewers or to freelance Fast & Fair reviewers. The unit of analysis was the review invitation, not the unique reviewer.

Reviewer invitation outcomes were classified as accepted, declined, or did not respond. For conventional peer review, “did not respond” was defined as no response within 10 days. For Fast & Fair, reviewers were expected to respond within the Fast & Fair response window, 1 working day. The distribution of invitation outcomes was compared between Fast & Fair and conventional peer review using a 2×3 chi-square test.

We also assessed review completion among reviewers who accepted invitations. A review was considered completed if the reviewer submitted a review report after accepting the invitation. Completion rates were calculated using accepted invitations as the denominator. Completion rates between Fast & Fair and conventional peer review were compared using Fisher’s exact test.

### Statistical analysis

All statistical analyses were performed using two-sided tests. Statistical significance was defined as p<0.05. For bounded or non-normally distributed continuous outcomes, groups were compared using Mann-Whitney U tests. For categorical outcomes with more than two categories, groups were compared using chi-square tests. For 2×2 categorical comparisons, Fisher’s exact test was used. No adjustment was made for multiple comparisons because each analysis addressed a distinct prespecified outcome: time to first decision with reviews, review quality, editorial decision profile, final acceptance rate, and reviewer commitment.

## ACKNOWLEDGEMENTS

We thank our academic editors, reviewers, and The Company of Biologists staff including Sue Chamberlain, Ania Crowther, Laura Tolhurst, Laura Mason, Andrea Kendall, Lindsey Cole, James Allison, and Rachel Hackett for their invaluable assistance. We thank Reinier Prosee for helping to recruit preLighters as reviewers. We also thank the Board of Directors at The Company of Biologists for their support, especially the members assigned to advise Biology Open: Jane Langdale, Steven Royle, Laura Machesky, Pleasantine Mill and Daniel St Johnston. We are grateful to Claire Moulton for advice throughout the experiment and Katherine Brown for comments on the manuscript.

